# Vascular dysregulation following SARS-CoV-2 infection involves integrin signalling through a VE-Cadherin mediated pathway

**DOI:** 10.1101/2022.03.15.484274

**Authors:** Danielle Nader, Steve Kerrigan

## Abstract

The vascular barrier is heavily injured following SARS-CoV-2 infection and contributes enormously to life-threatening complications in COVID-19. This endothelial dysfunction is associated with the phlogistic phenomenon of cytokine storms, thrombotic complications, abnormal coagulation, hypoxemia, and multiple organ failure. The mechanisms surrounding COVID-19 associated endotheliitis have been widely attributed to ACE2-mediated pathways. However, integrins have emerged as possible receptor candidates for SARS-CoV-2, and their complex intracellular signalling events are essential for maintaining endothelial homeostasis. Here, we showed that the spike protein of SARS-CoV-2 depends on its RGD motif to drive barrier dysregulation through hijacking integrin αVβ3. This triggers the redistribution and internalization of major junction protein VE-Cadherin which leads to the barrier disruption phenotype. Both extracellular and intracellular inhibitors of integrin αVβ3 prevented these effects, similarly to the RGD-cyclic peptide compound Cilengitide, which suggests that the spike protein – through its RGD motif – binds to αVβ3 and elicits vascular leakage events. These findings support integrins as an additional receptor for SARS-CoV-2, particularly as integrin engagement can elucidate many of the adverse endothelial dysfunction events that stem from COVID-19.

## Introduction

The interaction between the spike protein of SARS-coronavirus-2 (SARS-CoV-2) and endothelial cells has been widely demonstrated to be a critical driver in vascular dysregulation observed in COVID-19. We were the first to describe a pattern of impaired vascular functionality following SARS-CoV-2 infection, and theorized that the major endothelial adherens junction protein, VE-Cadherin, was involved^1^. A plethora of data now confirms this finding, where the disruption of junction proteins leads to reduced endothelial barrier integrity and subsequent monolayer permeability, elucidating the vast cardiovascular complications and septic shock experienced in severe COVID-19^2^. Although the canonical ACE2 receptor has been implicated in driving this reaction, another potential mechanism of action involves an integrin-mediated pathway. These heterodimeric transmembrane proteins are key regulators of haemostasis, angiogenesis, proliferation, and inflammation. Activated through binding an RGD-containing ligand, integrins can control downstream signalling transduction cascades that tether VE-Cadherin at the cell junctions through RhoGTPase cycling^3,4^. The spike protein contains an integrin-binding RGD motif that adheres to integrins αVβ3 and α5β1 on pulmonary epithelial cells and endothelial cells, where integrin antagonists Cilengitide and ATN-161 have demonstrated success in inhibiting this interaction *in vitro* and *in vivo*, thereby suggesting integrin-targeted therapeutics in COVID-19^1,5–8^. We aimed to identify the direct pathway that associates integrins with vascular dysregulation during SARS-CoV-2 infection, and whether targeting the spike protein RGD motif is sufficient to reduce this disease phenotype.

## Methods

### Cell and virus culture conditions

Primary-derived Human Aortic Endothelial Cells (HAoEC; Promocell C-12271) were maintained in Endothelial Cell Media MV (PromoCell) supplemented with 10,000 U/mL Penicillin and 100 mg/mL Streptomycin. Cells were subject to 10 dynes/cm^2^ of shear stress. Human 2019-nCoV strain 2019-nCoV/Italy-INMI1 was obtained from the European Virus Archive Global (Ref. no: 008V-03893) and experiments were carried out using a Multiplicity of Infection (MOI) of 0.4.

### ELISA

Interactions between recombinant spike proteins and integrins were performed as described previously^1^. Briefly, a 96-well microplate was coated with 25 ng of integrin αVβ3 (3050-AV, R&D Systems) overnight at 4°C and blocked in 5 % dry milk in 0.1 % Tween 20-PBS. Anti-αVβ3 mAB (MAB1876-Z, 1:100), anti-β3 mAB (sc-46655, 1:100), Cilengitide (0.0005-0.05 μM), GLPG-0187 (0.05-10 μM) and Spike proteins (40591-V08H41, 40591-V08H23, 50 nM) were added and washed. After incubation for 1 hour with AlexaFluor 405-labeled spike protein antibodies (FAB105403V, 1:100), absorbance was measured at 405 nm.

### Endothelial permeability and immunofluorescence assays

Endothelial barrier injury and VE-Cadherin expression was measured using transwell permeability assays and immunofluorescence as described previously^1^. Briefly, HAoEC were grown to confluence on top chambers of inserts and infected with SARS-CoV-2 for 24 hours. Cells were pre-treated for 30 minutes with Src and FAK inhibitors (SU6656, PF562271, 1 μM). Fluorescein isothiocyanate-dextran (250ug/mL, 40kDa, Sigma-Aldrich) was added to the chambers and fluorescent intensity measured at 490/520 nm wavelengths. Cells grown on glass slides were infected and stained using anti-VE-Cadherin mouse monoclonal IgG1 antibody, conjugated to AlexaFluor 488 (F-8 sc-9989, 1:100), overlaid onto Fluoroshield mounting medium (ab104139). To measure internal VE-Cadherin, cells were acid washed briefly following antibody staining. Cells were images using AxioObserver Z1 microscope.

### Western blot analysis of VE-Cadherin

HAoEC were grown to confluency, pre-treated with Cilengitide (0.0005 μM) and infected with SARS-CoV-2 for 24 hours. Scraped supernatants were collected in 150 μL RIPA buffer. Proteins (10 μg) were loaded into an SDS-PAGE and stained using anti-VE-Cadherin mouse antibodies (F-8, sc-9989, 1:200) followed by anti-mouse secondary antibodies (sc-525405, 1:5000). GAPDH (ab-8245) was used as a loading control.

## Results

### SARS-CoV-2 Variants of Concern recognize integrin αVβ3 through conserved RGD site

*In silico* modelling of the spike protein RGD site (403-405) located with the receptor binding domain (RBD) has predicted numerous points of contact with its putative integrin receptor, αVβ3^1^. It has been proposed that select mutations within SARS-CoV-2 variants of concern enable a more wide-open RBD, thereby increasing the likelihood of host-virus recognition^9^. We performed ELISA-based assays to confirm the direct binding between integrin αVβ3 and spike protein of wild-type, Delta (B.1.617.2), and Omicron (B.1.1.529), in the presence or absence of the cyclic RGD peptide compound Cilengitide and neutralising monoclonal antibodies which target the active site of the αVβ3 integrin or the β3 subunit. Cilengitide was successful in reducing the association between the integrin and spike protein, similarly to antibodies, revealing the interaction is likely RGD-dependent (Fig. 1A,B). Comparatively, other integrin antagonists have demonstrated efficacy in reducing spike protein adherence to integrins such as GLPG-0187, a broad-spectrum pan integrin inhibitor^8,10^. Although successful at high concentrations, GLPG-0187 was unable to maintain sufficient blocking between the proteins, unlike Cilengitide which remained effective even at subnanomolar levels (Fig. 1C).

**Fig 1.**
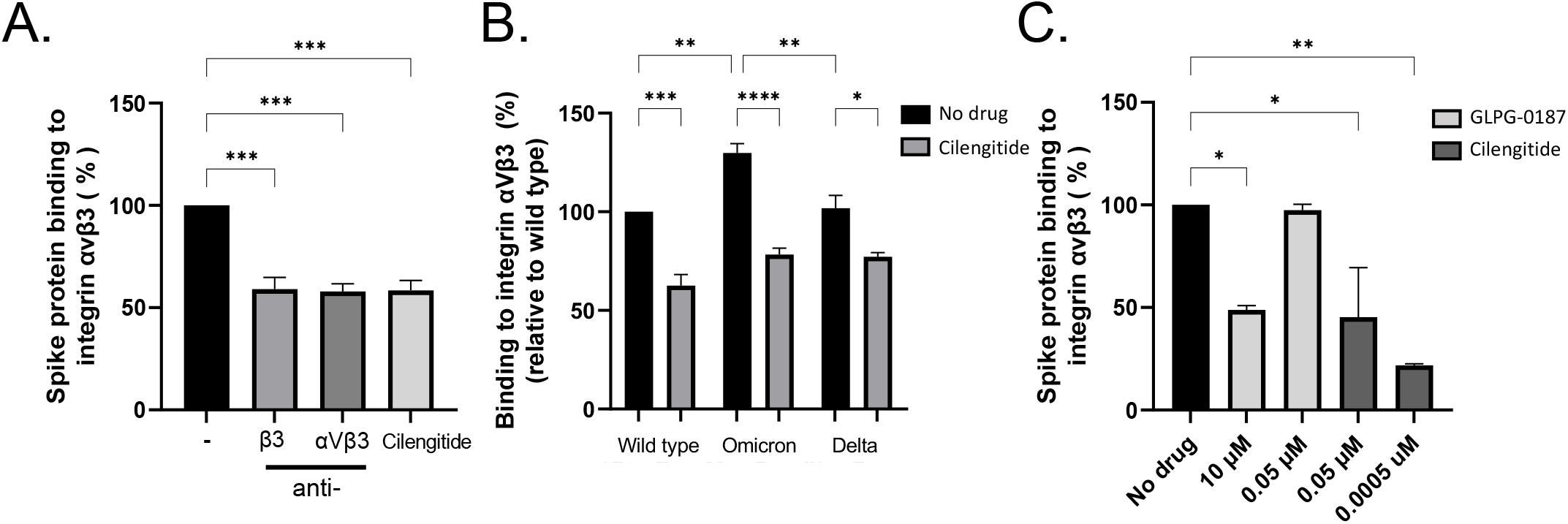
Wild type, Delta, and Omicron Spike proteins bind integrin αVβ3 through its RGD site. (A) Binding of SARS-CoV-2 spike protein to recombinant integrin αVβ3 in an ELISA-based assay. Integrin-blocking antibodies αVβ3 and β3 inhibit the interaction between recombinant spike protein and αVβ3, similarly to Cilengitide (0.0005 μM) (One-way ANOVA, ***P<0.001). (B) Effects of Cilengitide on variants of concern Delta and Omicron spike proteins binding to integrin αVβ3 (One-way ANOVA, *P<0.05, **P<0.01, ****P<0.0001). (C) Effects of broad-spectrum pan integrin inhibitor GLPG-0187 on spike protein binding to integrin αVβ3, compared to Cilengitide (One-way ANOVA). Values are mean ± S.E.M., n=3.

### Spike protein causes VE-Cadherin internalization which disrupts vascular permeability

Following spike protein engagement of integrins, reduced expression of some intercellular junction proteins (JAM-A and Connexin-43) has been detected in cerebral microvascular cells, alongside downregulation of the major adherens junction protein VE-Cadherin, which functions to mediate cell-cell adhesion^2^. Furthermore, evidence suggests SARS-CoV-2 spike protein directly induces this hyperpermeability through its RGD site, as treatment using the RGD peptide compound Cilengitide reduced inter-endothelial gaps and restored barrier function^1^. However, RGD-recognizing integrins are known to spatio-temporally coordinate the intracellular cycling of specific RhoGTPases to control VE-Cadherin without stimulating its downregulation^11^. To investigate this matter, we evaluated whether the endothelium could experience hyperpermeability through VE-Cadherin localisation during SARS-CoV-2 infection. Cell-surface or external VE-Cadherin levels were drastically reduced following viral infection for 24 hours, and blocking the RGD-binding site of integrins on host endothelial cells prevented this phenomenon (Fig. 2A,B). To measure non-surface bound VE-Cadherin, we utilised an acid-wash immunofluorescence staining protocol that enables the detection of internalised proteins. The amount of VE-Cadherin trafficked to intracellular compartments was significant in viral infected cells, revealing that intricate VE-Cadherin dynamics are likely involved in modulating endothelial permeability during COVID-19.

**Figure 2.**
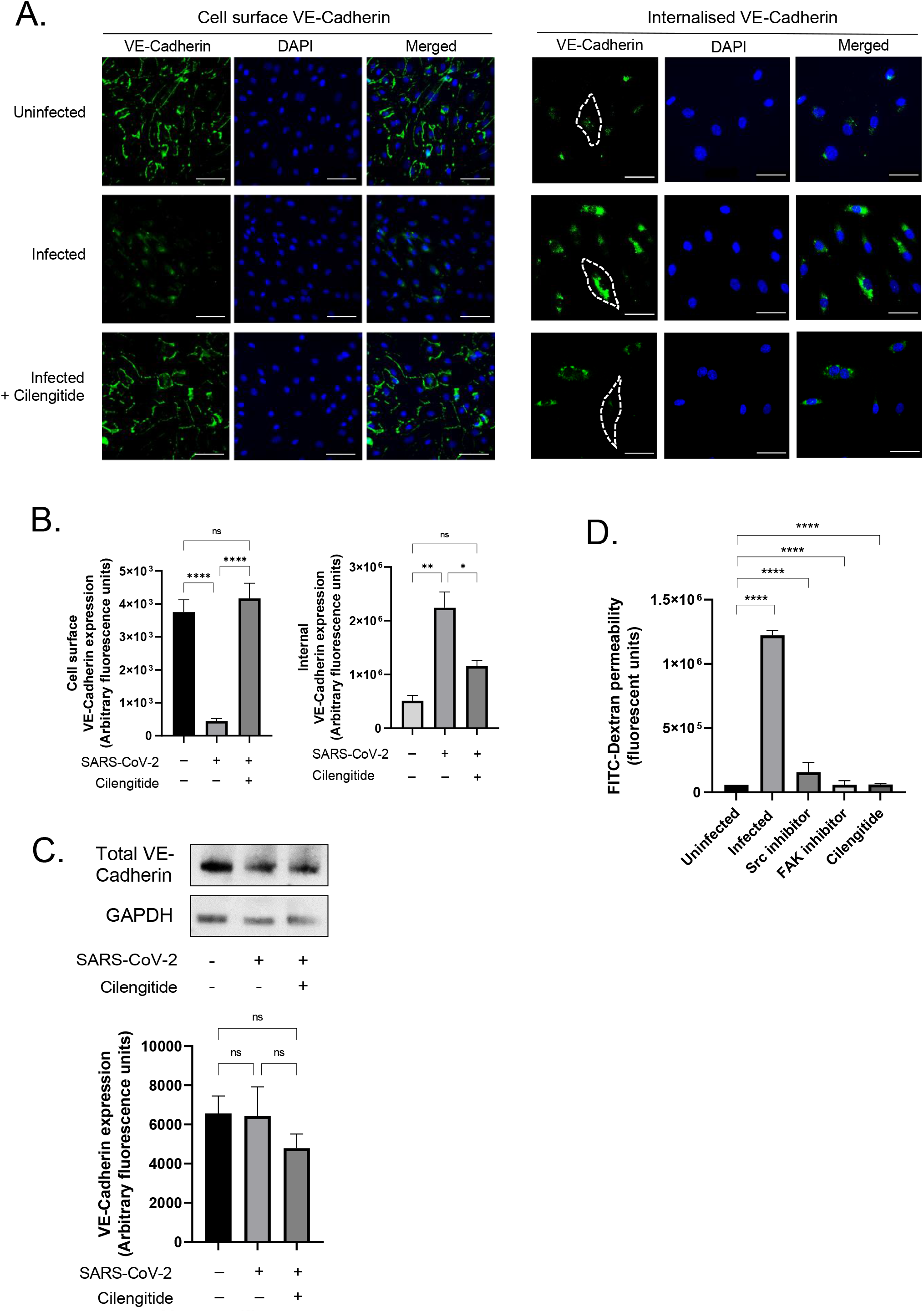
Vascular-Endothelial Cadherin is targeted by RGD site of SARS-CoV-2 spike protein to drive vascular dysfunction. (A) Immunofluorescence images of a confluent human endothelial cell monolayer stained with VE-Cadherin and DAPI, measured for either cell-surface or internal VE-Cadherin. Dotted lines represent the endothelial cell border as visualised using DIC. (B) Quantification of VE-Cadherin levels computed using ImageJ analysis, following background removal (One-way ANOVA). (C) Western blot analysis performed on total VE-Cadherin expression in healthy and SARS-CoV-2 infected endothelial cells. Treated cells were incubated with 0.0005 μM Cilengitide. Representative densitometry performed on western blot. (D) Transwell permeability assays measured endothelial barrier integrity over 24 hours of SARS-CoV-2 infection, in the presence of either Src and FAK inhibitors or Cilengitide. Values are mean ± S.E.M., n=3.

### Cilengitide reduces spike-induced endothelial dysregulation

Treating the infected cells with the αVβ3 integrin antagonist Cilengitide reduced the amount of internal VE-Cadherin to nearly uninfected basal levels (Fig. 2A,B). Our findings reveal that the spike protein binding to integrin αVβ3 directly triggers the integrin-mediated VE-Cadherin pathway in endothelial cells responsible for controlling vascular permeability. Moreover, the RGD site of the spike protein drives this pathway. Our data additionally revealed that overall VE-Cadherin levels were consistently stable in both healthy and infected vascular endothelial cells (Fig. 2C).

### Targeting intracellular integrin pathways can prevent vascular dysregulation following infection

Focal adhesion kinase (FAK) and Proto-oncogene tyrosine-protein kinase Src (Src) are non-receptor tyrosine kinases that localise to the integrin β tail, and are widely implicated in coordinating integrin signalling transduction in response to an external stimuli such as adhering to an RGD ligand. Since these proteins regulate the RhoGTPases that control VE-Cadherin internalisation, we investigated whether inhibiting these proteins associated with the integrin could also encourage similar events of vascular barrier protection as the integrin antagonist. Inhibition of FAK and Src prevented endothelial hyperpermeability in response to SARS-CoV-2 infection over 24 hours. Furthermore, this reduction was comparable to Cilengitide (Fig. 2D).

## Discussion

Clinical observations of viral endotheliitis, pulmonary thrombosis, hypoxia, oedema, and acute cardiac injury in patients with severe COVID-19 is indicative of a dysfunctional endothelial barrier, which establishes it as a vascular disease^12^. The relationship between SARS-CoV-2 spike protein and its host receptor ACE2 has been well defined, and a dual-receptor mechanism has been proposed with another cell surface receptor, integrins. In particular, integrins αVβ3 and α5β1 recognize the RGD motif uniquely expressed by SARS-CoV-2, which mediates infection of epithelial and endothelial cells *in vitro* and *in vivo*^1,5–8^. Subsequently, inter-endothelial junction weakening and hyperpermeability has been observed, which likely elucidates the pulmonary and cardiovascular complications in COVID-19^1,2,7^. Inhibiting spike protein attachment hinders this response^1,7^. Therefore we sought to describe the pathway correlating integrins to COVID-19 vascular dysregulation via VE-Cadherin.

Our work has identified the downstream signalling transduction cascade that links integrins directly to the observations of vasculopathy in COVID-19. Firstly, the spike proteins of Delta and Omicron SARS-CoV-2 variants of concern are still highly recognized by integrin αVβ3 as they both retain the RGD site. Both Cilengitide and integrin neutralising antibodies similarly blocked spike binding to integrins, revealing that this interaction is likely RGD-dependent. Other integrin antagonists such as GLPG-0187 have successfully displayed efficacy in reducing spike protein infection when used at high concentrations, confirming its involvement as a spike protein receptor^8^. However, when tested at similar concentrations to Cilengitide (0.05μM), it failed to prevent attachment. This may be due to the broad-spectrum activity of GLPG-0187 compared to the highly specific affinity Cilengitide has towards αVβ3 (IC50=0.58 nm). Similarly to the α5β1 antagonist ATN-161, Cilengitide has undergone clinical trials for the treatment of glioblastoma where it was greatly tolerated by patients due to its notable safety profile^13^. This has critical implications for COVID-19 treatment. Neutralizing antibodies recognize the ACE2 binding interface located on the spike protein surface (residues 437 – 507), and therefore mutations affecting receptor recognition often result in antibody evasion. Some *in vivo* and *in vitro* success has been observed using antibody cocktails to reduce viral load in SARS-CoV-2, particularly Etesevimab and Bamlanivimab combined therapy^14^. However, both antibodies were sensitive to mutations found in circulating variants of concern B.1.351 and B.1.617.2, and B.1.1.529 was partially or completely resistant to 100% of neutralizing monoclonal antibodies^15^. The RGD (403-405) motif is located within the spike receptor binding domain and is conserved across >99% of variants. As vaccine and immune-induced immunity is a key contributor to viral evolution, developing a compound that targets less immunodominant epitopes such as the RGD motif could be a more effective strategy against SARS-CoV-2 variants.

Our group was the first to identify that intercellular proteins were involved in SARS-CoV-2 pathogenesis and this likely corresponded to the hyperpermeability observed across the endothelium in COVID-19^1^. We previously identified the major adherens junction protein VE-Cadherin to be directly impacted during viral infection, where it was notably missing from its expected occupancy at the cell-cell contacts. When endothelial cells typically undergo angiogenesis and cell migration is required, the Rho GTPases Rac1 and RhoA tightly regulate stress fiber formation, whose spatially coordinated activation are triggered by integrins^11^. Subsequently, VE-Cadherin can undergo translocation into intracellular compartments via clathrin-mediated endocytosis, upon integrin activation and downstream transduction signalling involving the major focal adhesion proteins FAK and Src^3^. We propose that SARS-CoV-2 hijacks integrins via its RGD motif and controls its signalling cascade to command endothelial permeability. This ensures severe hypoxia and serum leakage, circulatory collapse and organ failure, which are key indicators of sepsis development. Critically, sepsis-related morbidity has been significantly attributed to COVID-19 deaths in both ICU and non-ICU patients. To explicate this matter, we evaluated the involvement of VE-Cadherin following spike protein infection. Although previous data suggests that VE-Cadherin was downregulated to some extent alongside other gap and tight junction proteins, here we showed that SARS-CoV-2 spike protein binding to integrin αVβ3 did not have any obvious effect on VE-Cadherin levels. However, infection dramatically altered VE-Cadherin organization by triggering its internalisation, which led to the dysfunctional barrier phenotype. Furthermore, the spike protein of SARS-CoV-2 has been demonstrated to signal the upregulation of RhoA in infected venous endothelial cells by downregulating Rac1, which promoted permeability and leakage^7^. Following integrin ligation, VE-Cadherin is known to coordinate with Rac1 to inhibit RhoA to regulate cell spreading^16^(Fig. 3). This is consistent with our own findings where we have found an association between αVβ3 and cell permeability, a process tightly controlled by VE-Cadherin.

**Figure 3.**
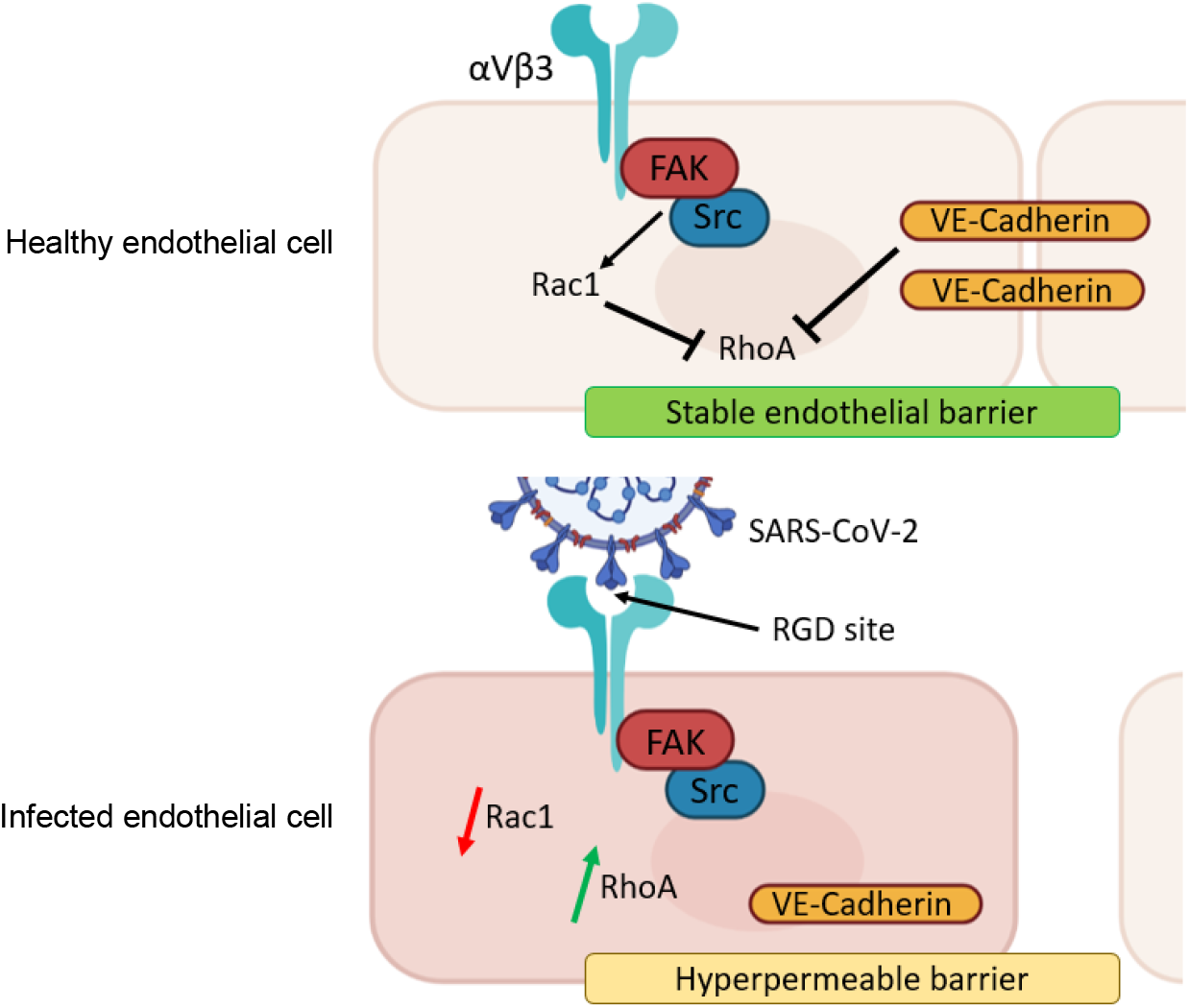
Schematic illustration depicting theorized intracellular signalling cascade following spike protein engagement with integrin. Top panel portrays healthy endothelial cell, where VE-Cadherin and Rac1 regulate and maintain low RhoA levels through FAK and Src signalling. Rac1 and RhoA signalling is tightly controlled via integrin engagement with an extracellular ligand. Bottom panel portrays infected endothelial cell, where persistent integrin activation leads to overactive FAK and Src activity, resulting in faulty cycling between RhoA and Rac1. RhoA levels rise, leading to cadherin phosphorylation via Src and FAK. Catenins, which confine VE-Cadherin at the endothelial junctions, cannot recognize phosphorylated VE-Cadherin which results it its internalization. This causes endothelial cells to pull apart and permeability to occur, promoting vascular leakage.

Additionally, several tyrosine sites across the cytoplasmic tail of VE-Cadherin undergo phosphorylation via FAK and Src proteins, and elevated phosphorylation of Y658 and Y731 accounts for the majority of barrier breakdowns^17^. Pharmacological inhibition of Src and its substrate FAK was effective in stabilizing VE-Cadherin at cell surface due to the significantly reduced endothelial permeability during SARS-CoV-2 infection. However, it has been reported that halting Src-mediated phosphorylation of VE-Cadherin is not especially sufficient to fully repair the endothelial barrier, suggesting a myriad of intracellular protein inhibitors would be required to suspend this occurrence^18^. Therefore, we suggest that eliminating the initial contact between viral and host protein – such as using integrin antagonists – would be more effective.

These early findings speculate that integrin engagement with the RGD-containing spike protein triggers a signalling cascade through FAK and Src, resulting in the downregulation of Rac1, increase in RhoA, and subsequent internalization of VE-Cadherin. Although some VE-Cadherin downregulation has been documented to occur, plasma membrane-associated VE-Cadherin can translocate into intracellular compartments following integrin activation through the RGD site of spike protein. As pools of endocytosed cadherins get recycled back to the plasma membrane, this could elucidate the process of vascular recovery in patients that do not experience severe COVID-19, although sepsis events can still take place^4,12^.

Altogether, we have shown that spike protein binding to integrin αVβ3 significantly impacts the integrity of the vascular barrier. Through its RGD site, SARS-CoV-2 effectively exploits downstream integrin signalling cascades to control permeability, which elucidates the dysfunctional vascular phenotype in COVID-19. A αVβ3 integrin inhibitor, Cilengitide, has displayed promising results in blocking this event. These early findings suggest that removing the initial signal that triggers integrin activation is capable of reducing the downstream signalling cascades that regulate cellular hyperpermeability in COVID-19. Evidently, endothelial cells are critical players during viral infection, and delineation of the mechanisms surrounding vascular integrity is required for the development of therapies to counteract the pathogenesis of SARS-CoV-2.

## Contributions

D.N. and S.K. wrote and edited the manuscript. D.N. performed the experiments. S.K. conceptualized the manuscript and provided critical review. All authors equally reviewed and approved the final manuscript.

## Acknowledgements

We would like to acknowledge the following individuals for their insight and conversations surrounding this topic: Gregory Bix, Timothy E. Gressett, Juan Pablo Robles, Prem Chapagain, and Tione Buranda.

## References

1 Nader, D., Fletcher, N., Curley, G. F. & Kerrigan, S. W. SARS-CoV-2 uses major endothelial integrin alphavbeta3 to cause vascular dysregulation in-vitro during COVID-19. PLoS One 16, e0253347, doi:10.1371/journal.pone.0253347 (2021).

2 Raghavan, S., Kenchappa, D. B. & Leo, M. D. SARS-CoV-2 Spike Protein Induces Degradation of Junctional Proteins That Maintain Endothelial Barrier Integrity. Front Cardiovasc Med 8, 687783, doi:10.3389/fcvm.2021.687783 (2021).

3 Li, R. et al. Vitronectin increases vascular permeability by promoting VE-cadherin internalization at cell junctions. PLoS One 7, e37195, doi:10.1371/journal.pone.0037195 (2012).

4 Xiao, K. et al. Mechanisms of VE-cadherin processing and degradation in microvascular endothelial cells. J Biol Chem 278, 19199–19208, doi:10.1074/jbc.M211746200 (2003).

5 Amruta, N. et al. In Vivo protection from SARS-CoV-2 infection by ATN-161 in k18-hACE2 transgenic mice. Life Sci 284, 119881, doi:10.1016/j.lfs.2021.119881 (2021).

6 Beddingfield, B. J. et al. The Integrin Binding Peptide, ATN-161, as a Novel Therapy for SARS-CoV-2 Infection. JACC Basic Transl Sci 6, 1–8, doi:10.1016/j.jacbts.2020.10.003 (2021).

7 Robles, J. P. et al. The spike protein of SARS-CoV-2 induces endothelial inflammation through integrin alpha5beta1 and NF-kappaB signalling. J Biol Chem, 101695, doi:10.1016/j.jbc.2022.101695 (2022).

8 Simons, P. et al. Integrin activation is an essential component of SARS-CoV-2 infection. Sci Rep 11, 20398, doi:10.1038/s41598-021-99893-7 (2021).

9 Md Lokman Hossen, P. B., Tej Sharma, Bernard Gerstman, Prem Chapagain. Significance of the RBD mutations in the SARS-CoV-2 Omicron: from spike opening to antibody escape and cell attachment. BioRxiv, doi: https://doi.org/10.1101/2022.01.21.477244 (2022).

10 Huntington, K. E. et al. Integrin/TGF-beta1 inhibitor GLPG-0187 blocks SARS-CoV-2 Delta and Omicron pseudovirus infection of airway epithelial cells which could attenuate disease severity. medRxiv, doi:10.1101/2022.01.02.22268641 (2022).

11 Pulous, F. E., Grimsley-Myers, C. M., Kansal, S., Kowalczyk, A. P. & Petrich, B. G. Talin-Dependent Integrin Activation Regulates VE-Cadherin Localization and Endothelial Cell Barrier Function. Circ Res 124, 891–903, doi:10.1161/CIRCRESAHA.118.314560 (2019).

12 Zhou, F. et al. Clinical course and risk factors for mortality of adult inpatients with COVID-19 in Wuhan, China: a retrospective cohort study. Lancet 395, 1054–1062, doi:10.1016/S0140-6736(20)30566-3 (2020).

13 Stupp, R. et al. Cilengitide combined with standard treatment for patients with newly diagnosed glioblastoma with methylated MGMT promoter (CENTRIC EORTC 26071-22072 study): a multicentre, randomised, open-label, phase 3 trial. Lancet Oncol 15, 1100–1108, doi:10.1016/S1470-2045(14)70379-1 (2014).

14 Gottlieb, R. L. et al. Effect of Bamlanivimab as Monotherapy or in Combination With Etesevimab on Viral Load in Patients With Mild to Moderate COVID-19: A Randomized Clinical Trial. JAMA 325, 632–644, doi:10.1001/jama.2021.0202 (2021).

15 Planas, D. et al. Considerable escape of SARS-CoV-2 Omicron to antibody neutralization. Nature 602, 671–675, doi:10.1038/s41586-021-04389-z (2022).

16 Nelson, C. M., Pirone, D. M., Tan, J. L. & Chen, C. S. Vascular endothelial-cadherin regulates cytoskeletal tension, cell spreading, and focal adhesions by stimulating RhoA. Mol Biol Cell 15, 2943–2953, doi:10.1091/mbc.e03-10-0745 (2004).

17 Potter, M. D., Barbero, S. & Cheresh, D. A. Tyrosine phosphorylation of VE-cadherin prevents binding of p120- and beta-catenin and maintains the cellular mesenchymal state. J Biol Chem 280, 31906–31912, doi:10.1074/jbc.M505568200 (2005).

18 Adam, A. P., Sharenko, A. L., Pumiglia, K. & Vincent, P. A. Src-induced tyrosine phosphorylation of VE-cadherin is not sufficient to decrease barrier function of endothelial monolayers. J Biol Chem 285, 7045–7055, doi:10.1074/jbc.M109.079277 (2010).

